# Cryo-EM structure determination of small proteins by nanobody-binding scaffolds (Legobodies)

**DOI:** 10.1101/2021.08.09.455715

**Authors:** Xudong Wu, Tom A. Rapoport

## Abstract

We describe a general method that allows structure determination of small proteins by single-particle cryo-electron microscopy (cryo-EM). The method is based on the availability of a target-binding nanobody, which is then rigidly attached to two scaffolds: (1) a Fab-fragment of an antibody directed against the nanobody, and (2) a nanobody-binding protein A fragment fused to maltose binding protein and Fab-binding domains. The overall ensemble of ∼120 kDa, called Legobody, does not perturb the nanobody-target interaction and facilitates particle alignment in cryo-EM image processing. The utility of the method is demonstrated for the KDEL receptor, a 23 kDa membrane protein, resulting in a map at 3.2Å overall resolution with density sufficient for *de novo* model building, and for the 22 kDa RBD of SARS-CoV2 spike protein, resulting in a map at 3.6 Å resolution that allows analysis of the binding interface to the nanobody. The Legobody approach thus overcomes the current size limitations of cryo-EM analysis.

## Introduction

Single-particle electron cryo-microscopy (cryo-EM) has become the method of choice for the determination of protein structures. Cryo-EM analysis has several advantages over X-ray crystallography or NMR ^1^, but the method becomes increasingly challenging for smaller proteins. Large molecules are relatively easy to identify in noisy low-dose images of vitrified samples and have sufficient contrast and features to determine their orientation and position for alignment and averaging. The structural analysis of small particles (∼100 kDa or less) is much more difficult. Without symmetry, they require optimal conditions, such as a highly homogeneous sample, rigid protein conformation, and random particle distribution in thin ice, conditions that are difficult to achieve with most samples ^2^. However, structure determination of small proteins is of great interest, as most proteins have sizes below 100 kDa and ∼50% are smaller than 50 kDa, including many membrane proteins and proteins of medical importance. It is thus a major goal in the field to expand the use of cryo-EM to the routine analysis of small proteins.

One approach to employ cryo-EM for small proteins is based on phase contrast methods, such as the use of Volta phase plates. This method has been used to determine the structure of streptavidin, a protein of 52 kDa, at 3.2 Å resolution ^3^. However, the structure of this protein could be determined even without phase plates ^4^, likely because streptavidin forms rigid tetramers and the particles display a near-perfect distribution in very thin ice, which greatly facilitates structural analysis.

An alternative strategy is to make the target protein larger, either by fusing it to another protein or by using a binding partner. In either case, high rigidity of the added scaffold itself and its rigid connection to the target protein are required to facilitate particle alignment and averaging in cryo-EM images.

The fusion approach has been tried with different scaffolds. For example, in a recent study, the BRIL domain was fused into a loop of a small GPCR protein by extending helices on both sides of the fusion point; the size of the scaffold was further increased by a Fab directed against the BRIL domain ^5^. However, this approach is limited to proteins containing suitable α-helices; their extension has to be customized for each new target to generate a rigid connection, which is difficult to achieve without prior knowledge of the target structure.

More promising is the use of a binding partner that can be selected with a screening platform, such as modified ankyrin repeat proteins (DARPins), Fab fragments of antibodies, or nanobodies. In recent studies, DARPins selected against GFP were grafted onto large scaffolds and used to visualize GFP by cryo-EM ^6, 7^. However, the intrinsic conformational heterogeneity of DARPins limits their potential to achieve high-resolution structures of small proteins ^7^, and so far only a few DARPins have been selected against membrane proteins. Fab fragments are mainly used in X-ray crystallography and only a few examples of their application for cryo-EM analysis have been reported ^8–10^, in part because the selection of appropriate Fabs is not trivial. In addition, the size of the Fabs (∼50 kDa) and the existence of a somewhat flexible hinge region between the two sub-domains still make structural analysis challenging.

Nanobodies, derived from single-chain antibodies of camelids, are also becoming popular as versatile binding partners of target proteins. Nanobodies have several attractive features. They form rigid structures that can bind to diverse shapes of target proteins, such as loops, convex surfaces, and cavities ^11^. They can be selected from immunized camelids or from large *in vitro* libraries displayed by phages, yeast cells, or on ribosomes ^11, 12^, and they can be produced in large quantities in a fairly short time. They often lock a protein into a fixed conformation, particularly in the case of membrane proteins, and have been used extensively to determine X-ray structures. The small size of nanobodies (12∼15 kDa) limits their direct application in cryo-EM, but the problem might be overcome if one could increase their size with the rigid attachment of a large scaffold. One reported attempt is to fuse a large protein into a loop of the nanobody, generating a “megabody” ^13^. However, the fusion did not result in a rigid connection of the scaffold to the protein of interest and it therefore did not sufficiently facilitate particle alignment, as indicated by the fact that the method has only been applied to the cryo-EM analysis of particles larger than 100 kDa.

Here, we describe a versatile method that allows cryo-EM analysis of even the smallest protein once a tightly binding nanobody is available. The size of the nanobody is increased to ∼120 kDa by two rigidly attached scaffolds. The overall design is reminiscent of a Lego construction, so we propose to call the scaffolds/nanobody ensemble “Legobody”. The utility of the Legobody method is demonstrated by structures of two small proteins (22 kDa and 23 kDa) that are asymmetric monomers and have a size well below the estimated limit for direct cryo-EM single-particle analysis (∼40 kDa) ^14^. The Legobody approach can easily be applied to any target protein and should greatly expand the use of cryo-EM single-particle analysis by overcoming the current size limitations.

## Results

### Generation of a nanobody-binding Fab

Our first nanobody-interacting scaffold is a Fab fragment of an antibody that is directed against a surface present in many nanobodies and not involved in target interaction. To generate such a Fab, we raised monoclonal antibodies in mice against a nanobody (Nb_0) that contains a framework sequence almost identical to that used in two libraries employed for rapid *in vitro* screening ^11, 12^. Amino acids in the antigen-interacting complementarity-determining regions (CDRs) of Nb_0 were chosen to minimize the immunogenicity of the antigen-interacting surface (see **Table S1** for sequence). To select for monoclonals that do not perturb nanobody-antigen interaction, hybridoma clones were screened with a complex of a nanobody against MBP (Nb_MBP) and its antigen MBP ^11^. After several rounds of selection, hybridoma clone 8D3 was obtained, which produced monoclonal antibodies that strongly bind to both Nb_0 and the Nb_MBP/MBP complex.

Although it is possible to directly use the antibodies secreted by clone 8D3 to generate Fab fragments, future applications are greatly facilitated if the Fabs can be made recombinantly. To this end, we first determined the DNA sequences of the regions coding for the variable regions of the light and heavy chains of clone 8D3. These sequences were then combined with the sequences coding for the constant regions of murine IgG1, and both genes were expressed together in HEK293 cells. Because the yield of Fabs was rather low, the constant regions of the light and heavy chains were replaced with those of human IgG. The resulting Fab_8D3 was expressed in HEK293 cells as a secreted protein and purified by Ni-NTA chromatography on the basis of a His-tag attached to the heavy chain. The yield is about 5∼8 mg from 1L of cell culture. Recombinantly purified Fab_8D3 forms a stable complex with nanobody Nb_0, as shown by co-migration of the proteins in size-exclusion chromatography (**Fig. 1a**).

**Fig. 1.**
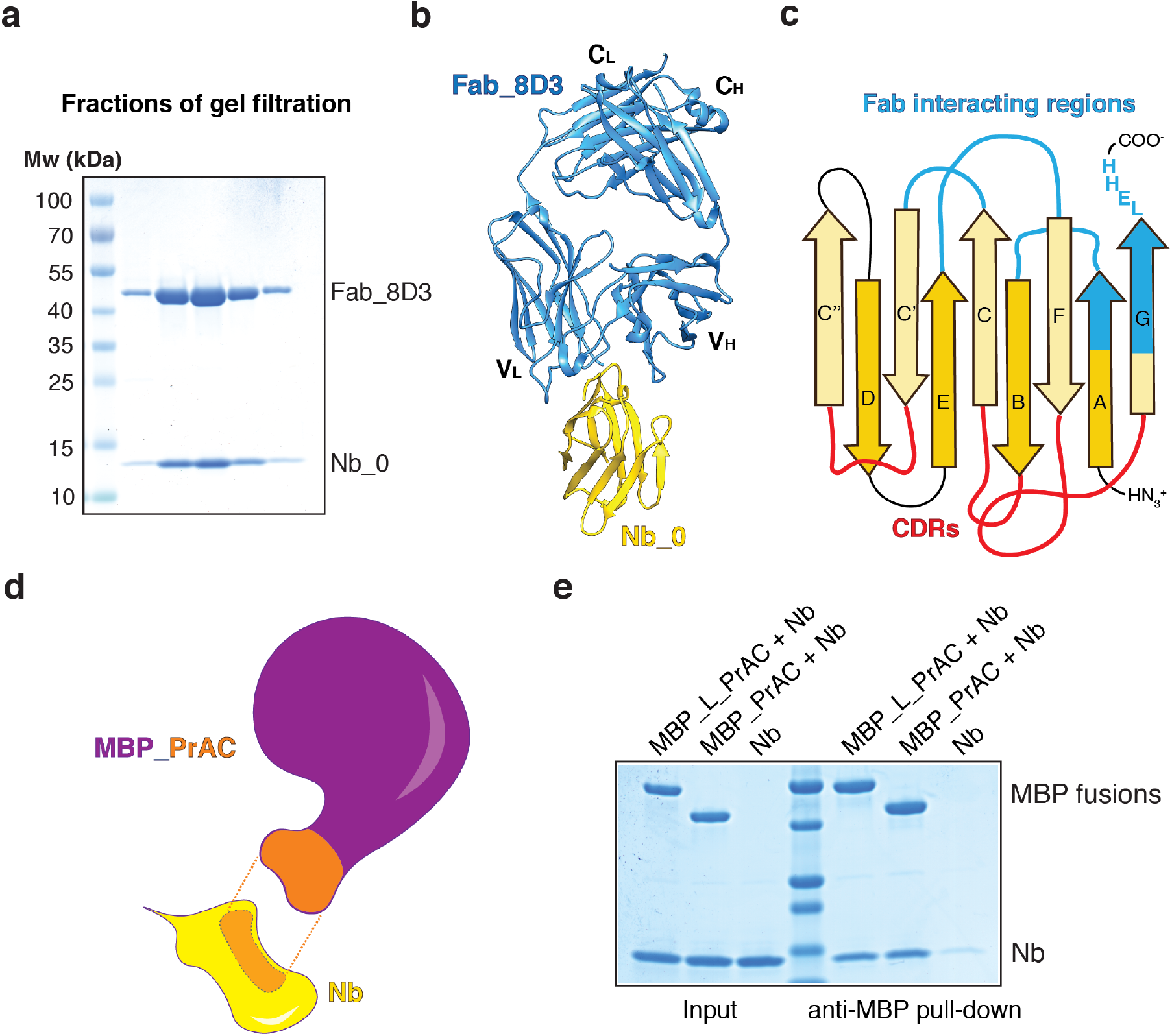
Generation of two nanobody-interacting scaffolds. **a**, Nanobody Nb_0 was mixed with Fab_8D3 and the complex was subjected to gel filtration. Fractions were analyzed by SDS-PAGE and Coomassie-blue staining. **b**, Crystal structure of the Nb_0/Fab_8D3 complex shown in cartoon representation. Nb_0 is colored in yellow and Fab_8D3 in blue. **c**, Scheme of the nanobody, with β-strands labeled according to convention. CDR loops are colored in red and regions interacting with the Fab in blue. **d**, Scheme of the interaction of nanobody (Nb; yellow) with the domain C of protein A (PrAC; orange) grafted onto maltose-binding protein (MBP; purple). The interacting regions between Nb and MBP_PrAC are shown in orange with dashed lines in between. **e**, PrAC fused to MBP through a flexible linker (MBP_L_PrAC) or grafted onto MBP (MBP_PrAC) was mixed with Nb. The mixture was incubated with MBP-interacting amylose resin, and the bound material analyzed by SDS-PAGE and Coomassie-blue staining. The input corresponds to 50% of the material used for the pull-down.

To identify the exact interaction surface, we determined a crystal structure of the complex of Fab_8D3 and Nb_0. After confirming that the crystals contained both components (**Fig. S1a**), a structure of the complex was determined at 1.8 Å resolution (**Fig. 1b; Table. S2**). As expected, Fab_8D3 binds to a surface of Nb_0 that is distal from the CDRs and contains conserved amino acids present in many nanobodies (**Fig. 1c; Fig. S1b**). The Fab-interacting amino acids of Nb_0 are located in loops between β-strands A and B, C and C’, E and F, as well as in segments of the β-strands A and G (**Fig. 1c**). Amino acids introduced by the cloning of the His-tag are also involved. The extensive interactions between the Fab and nanobody generate a rigid interface, a conclusion supported by the B-factor profile of the X-Ray structure (**Fig. S1c**).

### Generation of an MBP-based scaffold interacting with nanobodies

The second scaffold was developed on the basis of reports that protein A from *Staphylococcus aureus* can bind to nanobodies ^15^. Protein A contains five repeats of three-helical bundles (domains A, B, C, D and E). All these domains associate with the constant region of IgG antibodies, but also bind with different affinities to the variable region of the heavy chain of some antibodies (human V_H_3 family) ^16^, a region that is similar in sequence to the common framework of many nanobodies. Consistent with this sequence homology, protein A has been reported to interact with nanobodies in a similar way as with Fabs ^17^. To identify the strongest binding protein A domain, we fused domain D (PrAD) and the most divergent domains C and E (PrAC and PrAE) through a long, flexible linker to MBP (MBP_L_PrAC, MBP_L_PrAD and MBP_L_PrAE) and tested these fusions for their interaction with a nanobody. Co-elution of the proteins in size-exclusion chromatography showed that all three domains interact with the nanobody, but domain C forms the most stable complex (**Fig. S2a**).

Next, we grafted domain C of protein A (PrAC; ∼6 kDa) to MBP to generate a larger nanobody-binding partner. MBP is frequently used as an N-terminal fusion partner, as it can increase the solubility of its fusion partners. Although fusions can be designed as helical extensions of the C-terminal helix of MBP and have been extensively used in X-ray crystallography ^18^, such linkers are not rigid enough for cryo-EM analysis. To generate a more rigid connection between PrAC and MBP, we used the “shared helix” approach ^19^, applying it to a helix of domain C that is not involved in nanobody interaction (**Fig. S2b**). Residues on one side of this helix were mutated to those of MBP’s C-terminal helix that face the core of MBP. The resulting construct MBP_PrAC (**Fig. 1d**) could be expressed in *E. coli* and purified in large quantities. Like the MBP fusion of PrAC containing a flexible linker (MBP_L_PrAC), MBP_PrAC interacted with the nanobody in pull-down experiments (**Fig. 1e**).

### Legobody Assembly

The nanobody surfaces interacting with Fab_8D3 and MBP_PrAC are not overlapping (**Fig. 2a, b**), providing an opportunity to use both scaffolds at the same time. To increase the stability of the ensemble, we fused to the C-terminus of MBP_PrAC two Fab-binding domains. One is the domain D of protein A (PrAD), which has been shown to interact with the variable region of the heavy chain of Fabs ^20^. The other is protein G (PrG), which is known to strongly interact with the constant region of the heavy chain ^21^. All domains were connected by short linkers, generating a scaffold designated MBP_PrA/G (**Fig. 2a, b**). To allow the interaction of PrAD with Fab_8D3, some residues in the variable region of the heavy chain were mutated based on the crystal structure of a Fab/PrAD complex (PDB code 1DEE), generating Fab_8D3_2. Avidity effects increase the binding constants of both the MBP_PrAC and Fab scaffolds for the nanobody, making the overall assembly very stable.

**Fig. 2.**
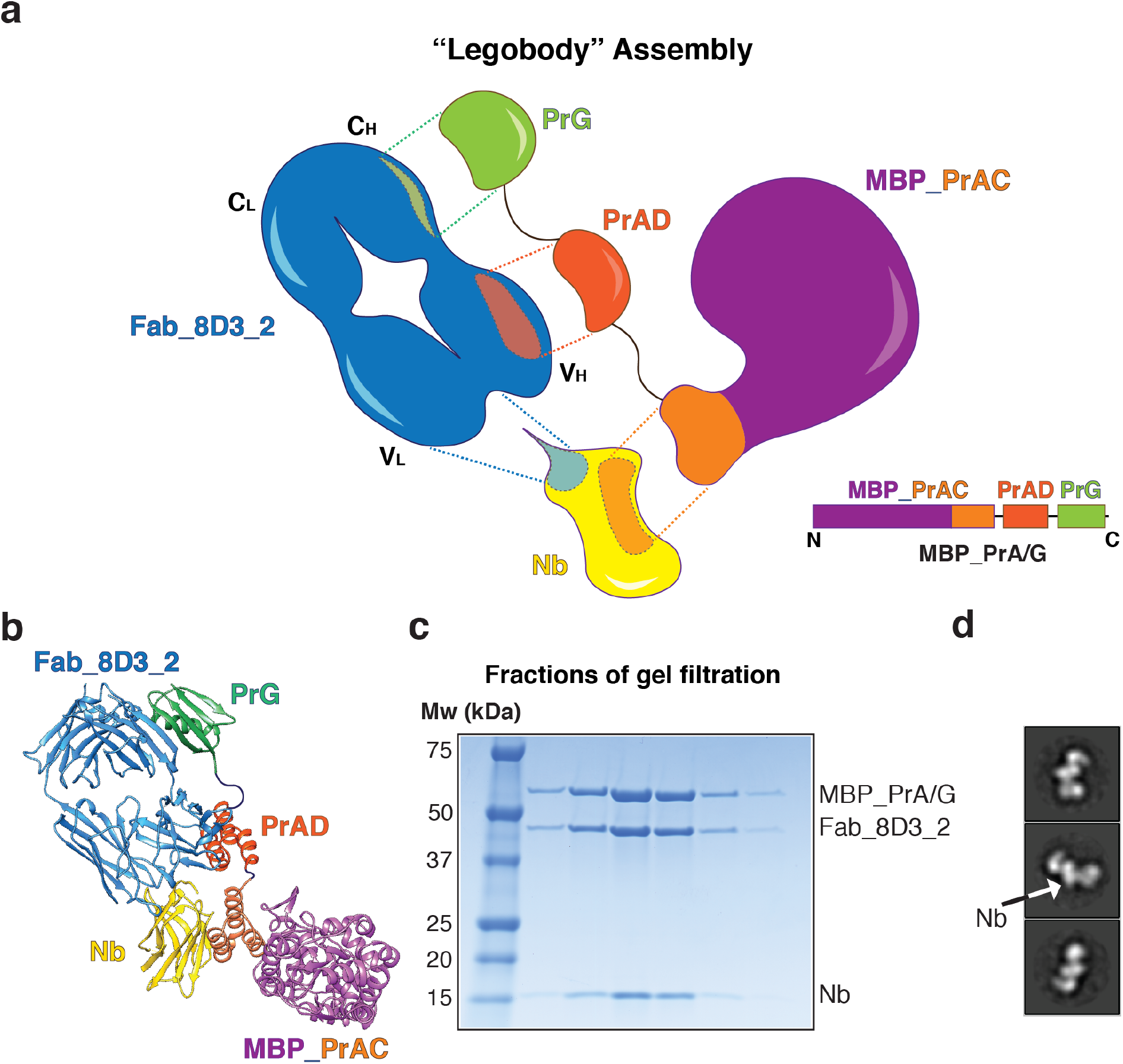
Assembly and purification of the Legobody. **a**, Scheme of the Legobody assembly. Fab_8D3-2, a derivative of Fab_8D3, and MBP_PrAC bind directly to the nanobody (Nb). The Fab-interacting D-domain of protein A (PrAD) and protein G (PrG) were fused sequentially to the C-terminus of MBP_PrAC via short linkers, generating MBP_PrA/G. Interacting surfaces are colored and highlighted by dashed lines. **b**, Model of the assembled Legobody in cartoon representation. The model was assembled from structures of Nb_0/Fab_8D3 (Fig. 1b), Fab/PrAD (PDB code 1DEE), and Fab/PrG (PDB code 1IGC), and by manually building a model for MBP_PrAC from the structures of MBP (PDB code 1ANF) and PrAC (PDB code 4NPD). **c**, Legobody assembled with a nanobody was subjected to gel filtration. Fractions were analyzed by SDS-PAGE and Coomassie-blue staining. **d**, Purified Legobody was analyzed by negative-stain EM. Shown are representative 2D class averages, with density for the nanobody highlighted by an arrow.

The complex between Fab_8D3_2 and MBP_PrA/G was assembled before adding a nanobody. All three components co-migrated in size-exclusion chromatography (**Fig. 2c**). Negative-stain EM showed strong structural features for the different parts of the Legobody (**Fig. 2d**), suggesting overall rigidity of the assembly, an assumption confirmed by subsequent cryo-EM analysis (see below). The assembly of the complex from individual pieces is reminiscent of a Lego construction, so we propose to call it a “Legobody”.

### Case study I: KDEL receptor (∼23 kDa)

To test the utility of the Legobody method, we first chose a small membrane protein, the KDEL receptor. This protein binds to the C-terminal KDEL sequence of luminal endoplasmic reticulum (ER) proteins that have escaped the ER, so that these proteins can be returned from the Golgi to the ER by vesicular transport ^22^. The KDEL receptor has seven trans-membrane segments and a molecular weight of only ∼23kDa. A crystal structure of the KDEL receptor in complex with a tightly binding nanobody has been reported ^23^.

To generate a sample suitable for cryo-EM analysis, we devised a protocol that should be applicable to many other challenging membrane proteins (**Fig. 3a**). The solubilized KDEL receptor (KDELR), tagged with a streptavidin-binding peptide, was first immobilized on streptavidin beads. We employed the detergent DMNG, as it resulted in a more homogeneous sample than DDM used for the crystal structure ^23^. The beads containing KDELR were then incubated with Legobody containing the reported nanobody against KDELR ^23^. Finally, to reduce aggregation during purification, the complex was reconstituted into nanodisc on the beads. After elution from the beads with biotin, the complex of KDELR, Legobody, and nanodisc was further purified by size-exclusion chromatography (**Fig. 3b**). Negative-stain EM showed features for the Legobody and nanodisc (**Fig. 3c**).

**Fig. 3.**
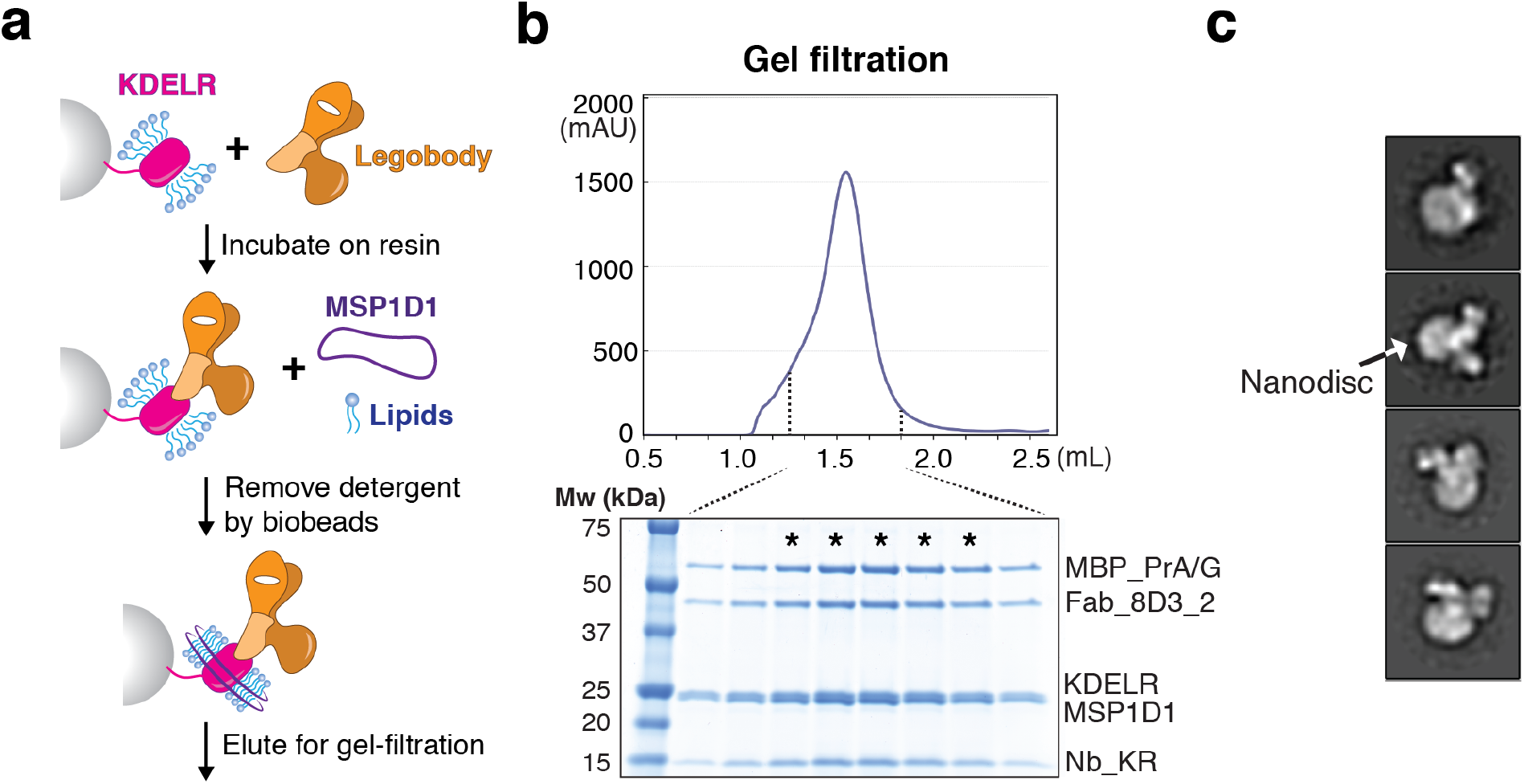
Purification of a KDEL receptor (KDELR)/Legobody complex. **a**, Scheme for the purification of the KDELR/Legobody complex. KDELR solubilized in detergent was immobilized on streptavidin beads. The resin was incubated with purified Legobody. Nanodisc-scaffolding protein MSP1D1 and lipids were added, and the reconstitution of nanodiscs initiated by the addition of Biobeads. After extensive detergent removal, the complex was eluted from the resin with biotin. **b**, The eluted complex of KDELR, Legobody, and nanodisc was subjected to gel-filtration and the elution of protein followed by the absorbance at 280 nm (upper panel). Fractions between the indicated dashed lines were analyzed by SDS-PAGE and Coomassie-blue staining (lower panel). Fractions indicated by a star were pooled and used for EM analysis. **c**, The purified complex was analyzed by negative-stain EM. Shown are representative 2D class averages, with density for the nanodisc highlighted by an arrow.

We next analyzed the complex by cryo-EM. When placed directly onto EM grids, the particles showed severe aggregation and strong preferred orientation, likely caused by denaturation of the molecules at the water-air interface. To alleviate this problem, surface lysine residues were modified with low molecular weight polyethylene glycol (PEG), a previously introduced method that makes the surface of proteins more hydrophilic and reduces their denaturation on the grids ^24^. Although the particles still showed some aggregation and preferred orientation (**Fig. S3**), the cryo-EM analysis was straightforward, as the size and unique shape of the Legobody greatly facilitated particle picking and 2D and 3D classification. The final 3D refinement of the selected particles resulted in a 3D reconstruction with an overall resolution of 3.2 Å and little directional differences in resolution (**Fig. S3; Table. S3**).

With the exception of the relatively flexible PrAD domain, all parts of the Legobody and KDELR had well-resolved structural features, allowing even the visualization of the maltose molecule bound to MBP (**Fig. 4a; Fig. S4a**). The local resolution ranged from 3.0 to ∼4.0 Å and showed that the interactions between the two scaffolds and the nanobody, as well as the connection between PrAC and MBP, are all rigid (**Fig. 4b**). Because the nanobody is tightly associated with the KDELR, and because the center of alignment of the Legobody is at the position of the nanobody, an excellent map was obtained for the KDELR (**Fig. 4c**). The local resolution of this part of the map ranged from 3.0 to ∼3.5 Å and all amino acid side chains of the KDELR were clearly visible. In addition, several bound phospholipid molecules could easily be identified. The cryo-EM structure of the KDEL/nanobody complex is almost identical to that obtained by X-ray crystallography (**Fig. S4b**) ^23^.

**Fig. 4.**
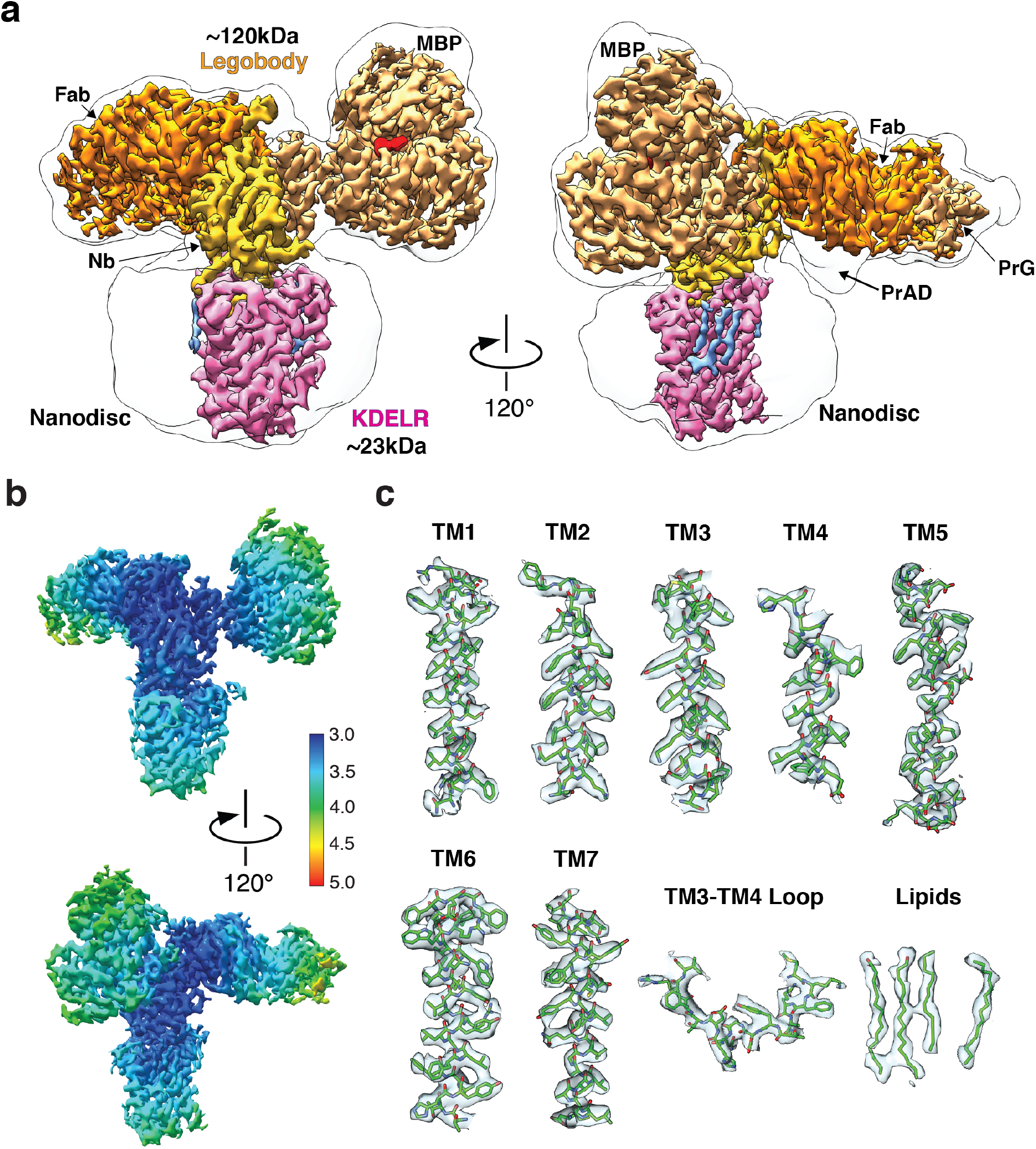
Cryo-EM structure of KDELR/Legobody complex. **a**, Cryo-EM map of the KDELR/Legobody complex in nanodiscs at 3.2Å resolution, shown in two views. The region of the KDELR is in pink and bound phospholipids are in light blue. The regions of the Legobody components are labeled and shown in different shades of yellow and orange. The density for MBP-bound maltose is shown in red. The solid line shows the outline of the cryo-EM map filtered to 10 Å, which allows the visualization of the nanodisc and flexible PrAD domain (right panel). **b**, Two views of the final map colored according to local resolution (see scale on the right). **c**, Map and fitted model for different segments of the KDELR and for bound phospholipids.

### Case study II: RBD of the SARS-CoV-2 spike protein (∼22 kDa)

Our second test protein for the Legobody method was the receptor-binding domain (RBD) of the SARS-CoV-2 spike protein. The RBD allows SARS-CoV-2 to bind to the ACE2 receptor and infect human cells ^25^. This interaction is of great medical interest, particularly during the current pandemic, and therefore many RBD-neutralizing nanobodies have been generated ^26^. The RBD has a molecular weight of only ∼22 kDa.

The RBD was expressed in HEK293 cells as a secreted protein and purified by Ni-NTA chromatography on the basis of an attached His-tag. The protein was mixed with the pre-assembled Legobody containing a reported nanobody against the RBD ^27^. The complex was further purified by size-exclusion chromatography (**Fig. 5a**) and analyzed by cryo-EM. After 2D and 3D classification, followed by 3D refinement, a map with an overall resolution of 3.6 Å was obtained, again with little directional differences in resolution (**Fig. 5b; Fig. S5a-e; Table. S3**). The local resolution ranged from ∼3.4 to ∼4.4 Å and showed good density for the central regions of the Legobody and target protein (**Fig. 5c**). Specifically, the RBD region showed good side-chain density for amino acids at the interface with the nanobody (**Fig. 5d**). The cryo-EM structure of the RBD/nanobody complex is almost identical to that obtained by X-ray crystallography (**Fig. S5f**) ^28^. Some polypeptide loops distal to the binding interface were invisible in the cryo-EM map (**Fig. S5f**). The biased particle orientation suggests that this may be caused by the partial denaturation of the RBD at the water/air interface, i.e. by issues unrelated to the Legobody method. The moderate preferred particle orientation observed with the KDELR and the somewhat stronger bias seen with the RBD are likely not caused by the Legobody itself, as the populated angles were different.

**Fig. 5.**
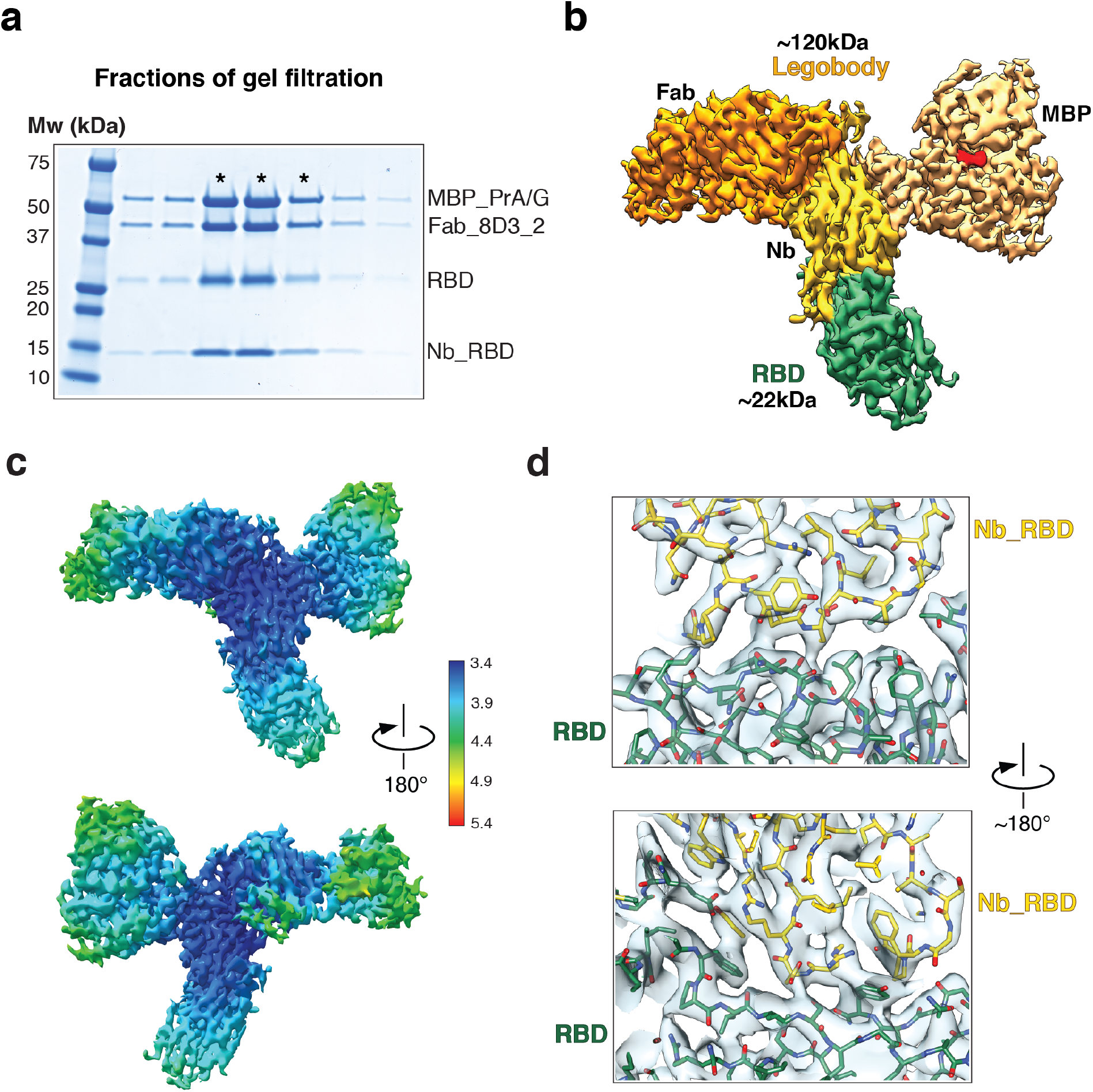
Cryo-EM structure of a complex of the RBD domain of the SARS-CoV-2 spike protein and Legobody. **a**, Purified RBD was mixed with Legobody containing a nanobody directed against the RBD (Nb_RBD). The sample was subjected to gel filtration and fractions were analyzed by SDS-PAGE and Coomassie-blue staining. Fractions labeled with a star were pooled and used for EM analysis. **b**, Cryo-EM map of the RBD/Legobody complex. The region of the RBD is shown in green. The regions of the Legobody components are labeled and shown in different shades of yellow and orange. The density for MBP-bound maltose is shown in red. **c**, Two views of the final map colored according to local resolution (see scale on the right). **d**, Map and fitted model for the interface between the RBD and nanobody.

### General applicability of the Legobody method

The Legobody method was developed using nanobodies that have a common framework similar to that used in two *in vitro* libraries ^11, 12^. However, the method can be applied to nanobodies that originally have a different framework, such as those often generated in alpaca. For example, a nanobody directed against the ALFAtag peptide, which is frequently used to purify or visualize fusion proteins ^29^, differs significantly in its framework sequence (sequence identity ∼75%) and would therefore not allow interactions with our scaffolds. Specifically, nine amino acid residues in the Fab- and PrAC-interacting regions were different from the ones in the common framework and thus mutated, resulting in Nb_ALFA that could be assembled into a Legobody (**Figs. S6a; S6b**). These mutations did not affect antigen binding, as GST-tagged ALFA peptide was able to pull-down the pre-assembled Legobody containing the modified nanobody (**Fig. S6b**). We therefore believe that all nanobodies can be used, regardless of whether they are obtained from *in vitro* libraries or from animal species.

## Discussion

Here we describe a general method that allows cryo-EM structures to be determined for small proteins. Our Legobody approach thus overcomes current limitations of cryo-EM analysis and greatly expands its use. The new method can be applied to any target protein once a tightly binding nanobody is available. The nanobody is assembled into a Legobody by the binding of two scaffolds, a Fab fragment and a MBP molecule to which domain C of protein A domain has been grafted (MBP_PrAC). All interactions were designed to be rigid. In addition, Fab-interacting domains were fused to MBP_PrAC to further solidify the complex. The Legobody has a characteristic shape, consisting of two lateral arms, formed by the two scaffolds, and a central lobe, contributed by the nanobody. The overall size (∼120 kDa) and shape of the Legobody, and the center of alignment at the position of the nanobody, greatly facilitate all steps of cryo-EM analysis, from particle picking, classifications, to final refinement. We demonstrate the utility of the Legobody method with two examples of small target proteins (KDELR (23kDa) and the RBD (22kDa) of the SARS-CoV-2 spike protein).

The membrane protein KDELR poses a particular challenge for cryo-EM analysis, as it is small, has no domains outside membrane, and no symmetry to facilitate particle alignment in EM images. The protein tends to aggregate during purification and on cryo-EM grids at the water-air interface of thin ice. To determine its structure, we not only used the Legobody approach, but also employed two other tricks, which likely are applicable to other challenging membrane proteins. First, we used a purification strategy, in which the KDELR/Legobody complex was incorporated into nanodisc while bound to beads (**Fig. 3a**). This strategy reduces aggregation of the receptor and increases its stability in solution. Second, before applying the sample to EM grids, we modified surface lysines with low-molecular weight PEG. We have used this protocol routinely for other proteins ^24, 30, 31^, and it often dramatically reduces particle aggregation and preferred particle orientation. Using standard cryo-EM data analysis, we were able to obtain a high-quality map for the KDELR, with density visible for all amino acid side chains. The map would have allowed straight-forward *de novo* model building.

The KDELR is representative of a large group of membrane proteins, which are of small size and pose similar challenges for cryo-EM analysis. Examples include G-protein coupled receptors, solute carrier transporters, and membrane-embedded enzymes, many of which are of great interest for drug development. The Legobody method now makes all these proteins accessible to cryo-EM analysis. Of course, the target proteins might adopt several physiologically relevant conformations and a nanobody might select only one of them.

The RBD of the SARS-CoV-2 spike protein also presents a challenge for cryo-EM analysis, as it is small and consists mainly of β-strands and extended polypeptide segments, which are more difficult to model into a map than α-helices. The map obtained with the Legobody approach was of good quality, especially at the RBD/nanobody interface, with side-chain density for all interacting amino acids. Because this interface is the region of medical interest, our results show that cryo-EM can be used to optimize RBD-neutralizing nanobodies, which may be important for the quick response to future virus pandemics. By comparison with X-ray crystallography, cryo-EM requires only small amounts of protein and can be performed in a significantly shorter time period.

A possible limitation of the Legobody method could arise from steric clashes between the scaffolds and targets. To test whether this is a serious problem, crystal structures of complexes of nanobody with small monomeric target proteins were aligned with the Legobody structure on the basis of the nanobody. Observed steric clashes are listed in **Table. S4**. For soluble proteins, only three examples were found in which the Fab or MBP_PrAC would clash with the target. For membrane proteins embedded into detergent micelles, nanodiscs, or lipid cubic phase, clashes would sometimes be caused by the PrAD domain or the MBP_PrAC scaffold. No clashes were observed for the Fab, but the number of available structures is too small to exclude their existence in other cases. Clashes with the PrAD domain can be avoided by deleting this domain and connecting MBP_PrAC directly to PrG via a suitable linker. In fact, this domain bound only weakly to its intended binding site on the Fab and should therefore be dispensable even in the original design. Tests for the compatibility of the scaffolds with the nanobody/target interaction are straightforward (similar to the pull-down experiments in **Fig. S6b**), which could also be used to quickly screen for nanobodies that are compatible with both scaffolds, avoiding nanobodies that would cause steric clashes.

Because of the modular design of the Legobody, only one of the two scaffolds can be attached to the nanobody. If only the MBP_PrAC scaffold is used, we recommend fusing the nanobody to the N-terminus of MBP_PrAC via a flexible linker. It should be noted that both scaffolds not only increase the size of the target, but also its solubility and monodispersibility. For example, the KDELR tends to aggregate during purification if either the Fab or the MBP_PrA/G scaffold are omitted.

The Legobody used in this study could easily be modified to further increase its molecular mass, stability, and rigidity, which would help to further improve the resolution of the cryo-EM maps. For example, the three-helix bundle of PrAC could be engineered to increase its binding affinity to the nanobody or it could be grafted onto other large proteins. In addition, fusions could be generated with protein L (ref. ^32^) or a modified version of protein M (ref. ^33^), which bind to the Fab at different sites than the fusion partners used in the current Legobody design. We believe that the current Legobodies and their possible variations will make cryo-EM structure determination of small proteins a routine method.

## Methods

### Purification of nanobodies

Genes for His-tagged or Strep II-tagged nanobodies were cloned into the pET 26b vector (Novagen). The expression and purification of all His-tagged nanobodies have been described previously ^12^. For immunization of mice, the His-tag was removed by treating purified nanobody Nb_0 with carboxypeptidase A (Sigma) and B (Roche) overnight at 4 °C. The treated nanobodies were passed through a Ni-NTA column (Thermo Fisher) and the flow-through fraction was further purified by size-exclusion chromatography on a S75 increase 10/300 GL column (Cytiva) in 25mM HEPES pH 7.4, 150mM NaCl, 5% glycerol. Strep II-tagged nanobody Nb_MBP was purified using streptactin resin (IBA). The beads were washed with 25mM HEPES pH 7.4, 150mM NaCl, and the protein was eluted in 25mM HEPES pH 7.4, 150mM NaCl, 2mM desthiobiotin. For screening of hybridoma cell clones, a complex of Strep II-tagged nanobody Nb_MBP and MBP was used. The eluted Strep II-tagged Nb_ MBP protein was mixed with separately purified MBP protein at a molar ratio of 3:1. The mixture was subjected to size-exclusion chromatography on a S200 increase 10/300 GL column (Cytiva) in 25mM HEPES pH 7.4, 150mM NaCl, 5% glycerol. The peak fractions of the complex were stored for future use.

### Generating antibodies against nanobodies

Monoclonal antibodies were generated by immunizing mice with purified tagless nanobody Nb_0 at the VGTI Monoclonal Core. Nanobody Nb_0 used for immunization and hybridoma screening also contained two mutations in the common framework (S7Y and S17Y) intended to increase the chances of selecting Fabs binding near that region, but the crystal structure showed that they were not involved in Fab binding. Hybridoma clones were screened under non-denaturing condition using the purified complex consisting of Strep II-tagged Nb_MBP and MBP. Antibodies secreted by clone 8D3 bound both Nb_0 and the Nb_MBP/MBP complex with high affinity. The 8D3 clone was expanded for further characterization. The sequences of the variable light (V_L_) and heavy (V_H_) chain regions of the monoclonal antibody were determined by Syd Labs.

### Recombinant expression of Fabs

To increase the yield of recombinant expression of the Fabs in HEK293 cells, the constant regions of the light and heavy chains were replaced by sequences from human Fabs (for sequences, see **Table. S1**). The resulting chimera genes for the light chain and the His-tagged heavy chain of the Fabs were separately cloned into the pCAGEN vector (a gift from Connie Cepko (Addgene plasmid #11160)) ^34^. For co-transfection of a 1L HEK293^freestyle^ (Thermo Fisher) culture, 0.5mg of both plasmids were incubated with 3mg of Linear PEI 25K (Polysciences) in 100ml of Opti-MEM (Thermo Fisher) medium at room temperature for 30 mins. The mixture was then added drop-wise into the medium containing HEK293^freestyle^ cells to reach a final cell density of 2 million/ml. The cells were cultured at 37 °C for 12∼16hrs before addition of 10mM sodium butyrate to boost expression. Medium containing secreted Fabs was harvested 48∼62hrs post-transfection. Purification of the Fabs was carried out as follows. Harvested medium free of cells was supplemented with 50mM TRIS pH 8.0, 200mM NaCl, 20mM imidazole and 1uM NiSO_4_. His-tagged Fabs were purified by Ni-NTA chromatography. The beads were washed extensively with 25mM HEPES pH 7.4, 200mM NaCl, 20mM imidazole. Fabs were eluted in 25mM HEPES pH 7.4, 200mM NaCl, 300mM imidazole. Eluted Fabs were concentrated and buffer-exchanged into 25mM HEPES pH 7.4, 150mM NaCl, 5% glycerol. Aliquots were snap-frozen for future use.

Based on reports that Fabs interact preferentially with domain D of protein A ^16^ and on a crystal structure of a complex of Fab and this domain (PDB code 1DEE), we introduced several mutations in the variable region of the heavy chain of the partially humanized Fab_8D3 (see above), resulting in Fab_8D3_2 (mutations: G16K, R18L, K19R, I58K, F80Y, T84N; the sequence is shown in **Table. S1**). Fab_8D3_2 was purified in the same way as the original Fab_8D3.

### Determination of a crystal structure of the Nb_0/Fab_8D3 complex

Purified His-tagged Fab_8D3 was mixed with purified His-tagged Nb_0 nanobody at a molar ratio of 1:3. The mixture was treated with carboxypeptidase A (Sigma) overnight at 4 °C to remove the His-tags. The sample was subjected to size-exclusion chromatography on a S200 increase 10/300 GL column in 25mM HEPES pH 7.4, 150mM NaCl. The peak fractions of the complex were pooled and concentrated to 10mg/ml and used to set up crystal screens.

Purified Nb_0/Fab_8D3 complex (0.2ul of a 10mg/ml solution) was mixed with 0.2ul of mother liquor containing 0.2M ammonium formate, 20-22% w/v PEG 3,350 using a Mosquito robot (TTP Labtech). Crystals were grown at 4°C with the hanging drop method over a reservoir of 100ul mother liquor and reached full size in about two weeks. Crystals were cryo-protected before harvest in a solution containing mother liquor supplemented with 25mM HEPES 7.5 and 18% ethylene glycol. X-ray diffraction data were collected on the 24-ID-E beamline at the Advanced Photon Source (APS). Initial phases were obtained by Molecular Replacement using crystal structures of Fabs and nanobodies with similar amino acid sequences as search models. In both search models, the CDR regions were removed.

### Purification of MBP fusions

The sequences of all MBP fusion proteins are given in **Table. S1**. Based on crystal structures and modeling, we predicted residue Ala405 of MBP_PrA/G-His6 to be close to the Fab and therefore mutated it to histidine to boost the interaction. All variants of MBP fusion proteins were purified as follows. The genes were cloned into the pET28b vector (Novagen) with either an N- or C-terminal His6 tag. The expression was induced by addition of 1mM IPTG for 4hrs at 37 °C. The cells were lysed by sonication in 25mM HEPES pH 7.4, 400mM NaCl, 20mM imidazole. The proteins were purified by Ni-NTA chromatography using lysis buffer as washing buffer. After elution with imidazole, proteins were subjected to size-exclusion chromatography on a S200 increase 10/300 GL column in 25mM HEPES pH 7.4, 150mM NaCl, 5% glycerol. The peak fractions were stored for future use.

### Purification of GST fused ALFA peptide

GST fused ALFA peptide was purified as follows. The gene for the ALFA peptide was cloned into the pGEX6p1 vector (Cytiva). The expression was induced by addition of 1mM IPTG for 5hrs at 30 °C. The cells were lysed by sonication in 25mM HEPES pH 7.4, 400mM NaCl. The proteins were purified by GST resins using lysis buffer as washing buffer. After elution with reduced glutathione (GSH), proteins were subjected to size-exclusion chromatography on a S200 increase 10/300 GL column in 25mM HEPES pH 7.4, 150mM NaCl, 5% glycerol. The peak fractions were stored for future use.

### Purification of Legobodies

Legobodies were assembled by first incubating purified MBP_PrA/G with Fab_8D3_2 at a molar ratio of 1: 1.1 in 25mM HEPES pH 7.4, 150mM NaCl. Then, the mixture was incubated with a chosen nanobody added at a 3-fold molar excess over MBP_PrA/G. The sample was applied to an amylose resin and the complex was eluted with 25mM HEPES pH 7.4, 150mM NaCl, 20mM maltose. The Legobodies were further purified by size-exclusion chromatography on a S200 increase 10/300 GL column in 25mM HEPES pH 7.4, 150mM NaCl, 5% glycerol, 2mM maltose. The peak fractions of the complexes were concentrated and stored for future use.

### Purification of a complex of KDEL-receptor (KDELR) and Legobody

The codon-optimized gene for the full-length KDELR with a SBP tag at its C-terminus was cloned into the pRS425-Gal1 vector (ATCC® 87331) ^35^. The expression of the receptor and preparations of the membrane fractions were carried out as previously described ^24^. Membranes from 15g of INVSc1 (Invitrogen) cells expressing the receptor were solubilized in 30ml of 25mM HEPES pH 7.4, 400mM NaCl, 1% DMNG (Anatrace) for 1hr. After removing insoluble material by ultra-centrifugation, the lysate was incubated with 250ul streptavidin resin (Thermo Fisher) for 1.5hr. The beads were collected and an excess of purified Legobody was added to the bound KDEL receptor to promote complex formation on the resin. After 1hr of incubation, the resin was washed with 8 column volumes of 25mM HEPES pH 7.4, 150mM NaCl, 2mM maltose, 0.02% DMNG. Nanodiscs were assembled on the resin by adding 1.25mM lipids (POPC/DOPE (Avanti) at a 4:1 ratio in DDM (Anatrace)) and 25uM nanodisc-scaffolding protein MSP1D1 in 700ul of washing buffer. After 30mins of incubation, the detergents were removed by the addition of two aliquots of 40mg of Biobeads and overnight incubation. The next day, the streptavidin resins were separated from Bio-Beads SM-2 (BioRad), taking advantage of their different rates of sedimentation by gravity. The streptavidin resins were washed by 25mM HEPES pH 7.4, 150mM NaCl, 2mM maltose and bound material was eluted with biotin. The KDEL receptor/Legobody complex in nanodisc was then purified by size-exclusion chromatography on a S200 increase 5/150 GL column in 25mM HEPES pH 7.4, 150mM NaCl, 2mM maltose. The peak fractions of the complex were concentrated, snap-frozen and stored for cryo-EM analysis.

### Purification of a complex of the SARS-CoV-2 spike RBD domain and Legobody

The codon-optimized gene for the RBD domain (residues 334-526) of SARS-CoV-2 spike protein with an N-terminal Flag tag and a C-terminal His8 tag was cloned into the pCAGEN vector. The RBD was expressed and purified in the same way as the Fabs. After elution from Ni-NTA beads, the protein was subjected to size-exclusion chromatography on a S75 increase 10/300 GL column in 25mM HEPES pH 7.4, 150mM NaCl. Peak fractions were mixed with Legobody at a molar ratio of 3:1. The mixture was subjected to size-exclusion chromatography on a S200 increase 5/150 GL column in 25mM HEPES pH 7.4, 150mM NaCl, 2mM maltose. The peak fractions of the complex were concentrated, snap-frozen, and stored for cryo-EM analysis.

### Cryo-EM Sample Preparation and Data Acquisition

The KDELR/Legobody complex at 0.8mg/ml was PEGylated by incubation with MS(PEG)12 methyl-PEG-NHS-ester (Thermo Fisher) at a 1:40 molar ratio for 2 hrs on ice to reduce preferred particle orientation on the grids. The chosen ratio allows a maximum of 1/3 of the total lysines to be modified, which minimizes effects of PEG modification on the stability of the complex. The PEGylated sample was then applied to a glow-discharged quantifoil gold grid (1.2/1.3, 400 mesh). The grids were blotted for 6∼7 s at 100 % humidity and plunge-frozen in liquid ethane using a Vitrobot Mark IV instrument (Thermo Fisher). The RBD/Legobody complex at 2.5mg/ml were incubated with MS(PEG)12 methyl-PEG-NHS-ester (Thermo Fisher) at a 1:25∼28 molar ratio for 2 hrs on ice. Right before plunge freezing, the PEGylated samples were diluted, using the gel-filtration buffers supplemented with detergent IGEPAL^®^ CA-630 (Sigma), so that the final protein and detergent concentrations were 1.2 mg/ml and 0.005%, respectively. The grids were frozen in the same way as described for the KDELR/Legobody sample.

Cryo-EM data for all samples were collected on a Titan Krios electron microscope (FEI) operated at 300 kV and equipped with a K3 direct electron detector (Gatan) at Harvard Cryo-EM Center for Structural Biology. A Gatan Imaging filter with a slit width of 25 eV was used to remove inelastically scattered electrons. All cryo-EM movies were recorded in counting-mode using SerialEM. For the KDELR/Legobody sample, the nominal magnification of 81,000x corresponds to a calibrated pixel size of 1.06 Å on the specimen. The exposure rate was 23.38 electrons/Å^2^/s. The total exposure time was 2.2 s, resulting a total electron exposure of 51.44 electrons/Å^2^, fractionated into 50 frames (44 ms per frame). For the RBD/Legobody sample, the calibrated pixel size was 1.06 Å. The exposure rate was 23.3 electrons/Å^2^/s. The total exposure time was 2.164 s, resulting a total electron exposure of 50.42 electrons/Å^2^, fractionated into 50 frames (44 ms per frame). The defocus range for both samples was between -1.0 and -2.6 µm.

### Cryo-EM Image Processing

For the KDELR/Legobody complex, dose-fractionated movies were subjected to motion correction using the program MotionCor2 ^36^ with dose-weighting. The program CtfFind4 ^37^ was used to estimate defocus values of the summed images from all movie frames. During data collection, the particles (close to 1 million) picked by YOLO ^38^ by “on-the fly” analysis using an automatic workflow at Harvard Medical School. The particles were then subjected to 2D classification (T2, 80 classes, 30 iterations) in Relion 3.1 ^39^. For 3D classification, an initial model was generated ab initio in Relion 3.1. After one round of 3D classification (T4, 5 classes, 50 iterations), there was only one class with clear protein secondary structure features. Particles of this class were selected for 3D refinement, resulting an initial reconstruction at 3.8 Å overall nominal resolution. This initial 3D reconstruction was used as 3D template to perform autopick in Relion 3.1 on the entire dataset, resulting in 2,532,161 particles. After 2D classifications (T2, 100 classes, 30 iterations), re-picked particles were “seeded” with particles used in the previous 3D refinement for 3D classification. After 3D classification (T4, 5 classes, 35 iterations), only the class showing clear protein secondary structure features of the whole complex was selected. After removing duplicates, the particles were subjected to 3D refinement, followed by polishing, CTF refinement, and another round of 3D refinement. Local 3D classification without image alignment (T20, 5 classes, 25 iterations) was performed using a mask including only the nanobody and KDELR. 246,878 particles were finally selected for 3D refinement using a mask excluding the nanodisc and the more flexible D domain of protein A.

For the RBD/Legobody complex, data analysis was performed in a similar way, except that particles from the “on-the-fly” analysis were not refined. Local resolution calculation and map sharpening were both performed in Relion 3.1. All reported resolutions are based on gold-standard refinement procedures and the FSC=0.143 criterion. Histograms of Directional FSC curves and Sphericity values were calculated with the 3DFSC server ^40^.

### Model Building

All model building was done in Coot. For the crystal structure of Nb_0/Fab_8D3, the initial phases were obtained by molecular replacement using the Phaser module in Phenix ^41^. The search models contained a nanobody, as well as the variable and constant regions of a Fab of similar framework sequence. After obtaining an initial density map, the model was refined with rigid bodies and then modified manually. The model was further refined using the Phenix.refine module with simulated annealing, XYZ, TLS, and individual B-factors. For the cryo-EM structures, initial models were based on the crystal structure of the Nb_0/Fab_8D3 complex, modified to account for the mutations in Fab_8D3_2, and the crystal structures of the KDEL receptor (6I6J), RBD (7KGJ), the manually grafted MBP_PrAC (1ANF and 4NPD), the D domain of Protein A (1DEE), and protein G (1IGC). These structures were docked into the maps and manually modified based on the cryo-EM density map. Models were then refined using the Phenix Real-space refinement module with minimization_global, local_grid_search, and ADP. For all refinements, secondary restraints, model restraints, and Ramachandran restraints were used.

## Acknowledgements

We thank S. Sterling, R. Walsh and M. Mayer at the Harvard Cryo-EM Center for Structural Biology for help in microscope operation and data collection. The X-ray structural work is based on research conducted at the Northeastern Collaborative Access Team beamlines, which are funded by the National Institute of General Medical Sciences from the National Institutes of Health (P30 GM124165). The Eiger 16M detector on 24-ID-E is funded by a NIH-ORIP HEI grant (S10OD021527). This research used resources of the Advanced Photon Source, a U.S. Department of Energy (DOE) Office of Science User Facility operated for the DOE Office of Science by Argonne National Laboratory under contract no. DE-AC02-06CH11357. We thank C. Bahl for initial efforts to use Rosetta for the design of fusions between MBP and Protein A domains, T-C. Hu for help with the generation of Fabs, the SBGrid team for software and workstation support, and Maofu Liao and Susan Shao for comments on the manuscript.

## Author contributions

X.W. conceived of ideas, designed and performed all the experiments, and analyzed the data. T.A.R. supervised the project. X.W. and T.A.R. wrote the manuscript.

## Data and materials availability

The coordinates of the atomic models of Nb_0/Fab_8D3, KDELR/Legobody and RBD/Legobody have been deposited in the Protein Data Bank with accession codes xxxx, xxxx and xxxx, respectively. The cryo-EM maps of the KDELR/Legobody and RBD/Legobody have been deposited with accession codes xxxx and xxxx, respectively. All other data and reagents are available on reasonable request from the corresponding authors.

## SUPPLEMENTARY MATERIALS

**Fig. S1.**
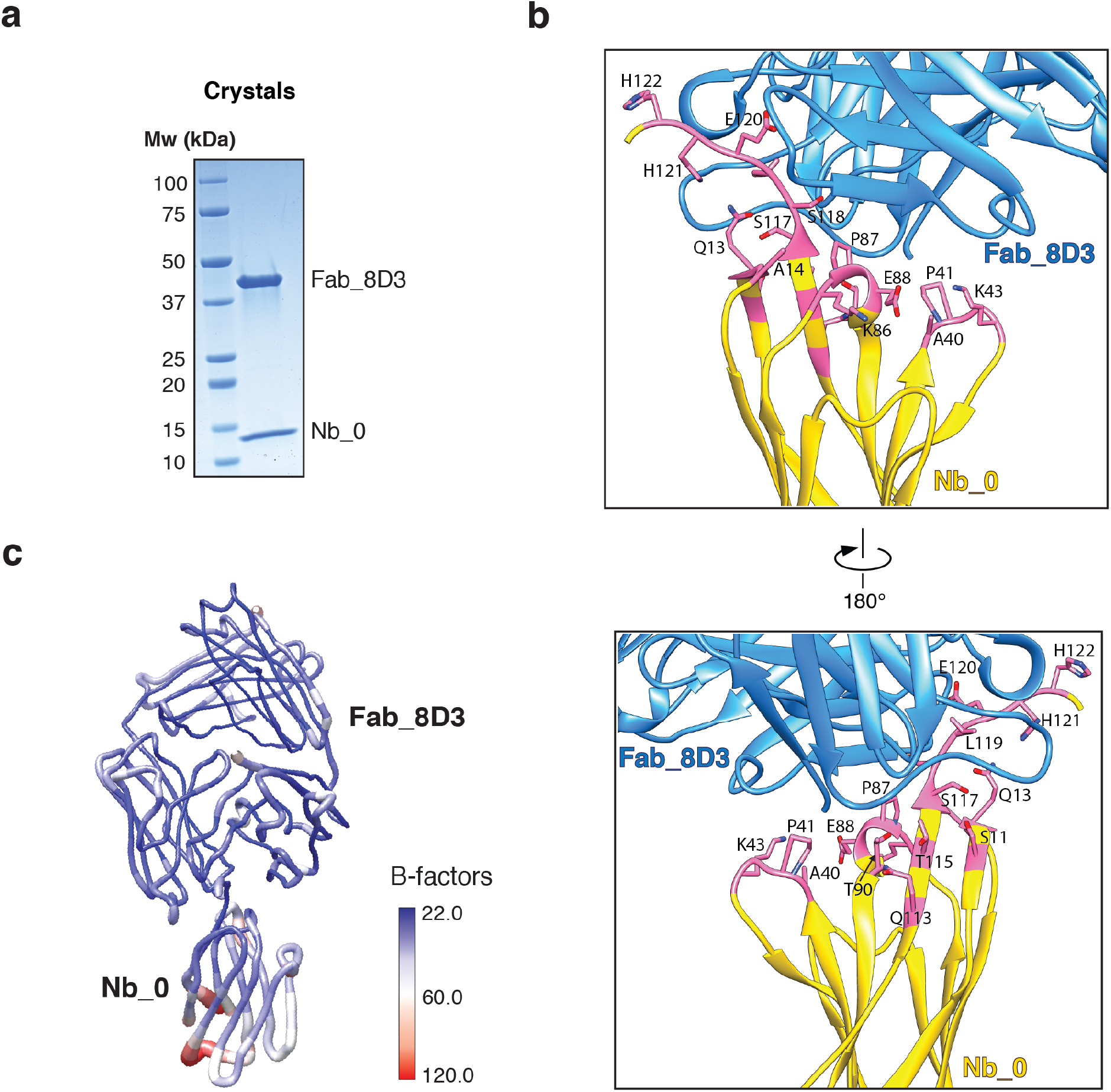
Crystal structure of a nanobody/Fab complex (Nb_0/Fab_8D3). **a**, Crystals contain both Nb_0 and Fab_8D3. Multiple crystals were pooled and analyzed by SDS-PAGE followed by Coomassie-blue staining. **b**, Two views of the interacting region of Nb_0 and Fab_8D3. Residues of Nb_0 involved in the interaction are colored in pink and labeled. The surface area buried at the binding interface is 898 Å^2^. **c**, Model of the Nb_0/Fab_8D3 complex in “worms” representation, colored according to the average B-factor of amino acid residues (scale on the right).

**Fig. S2.**
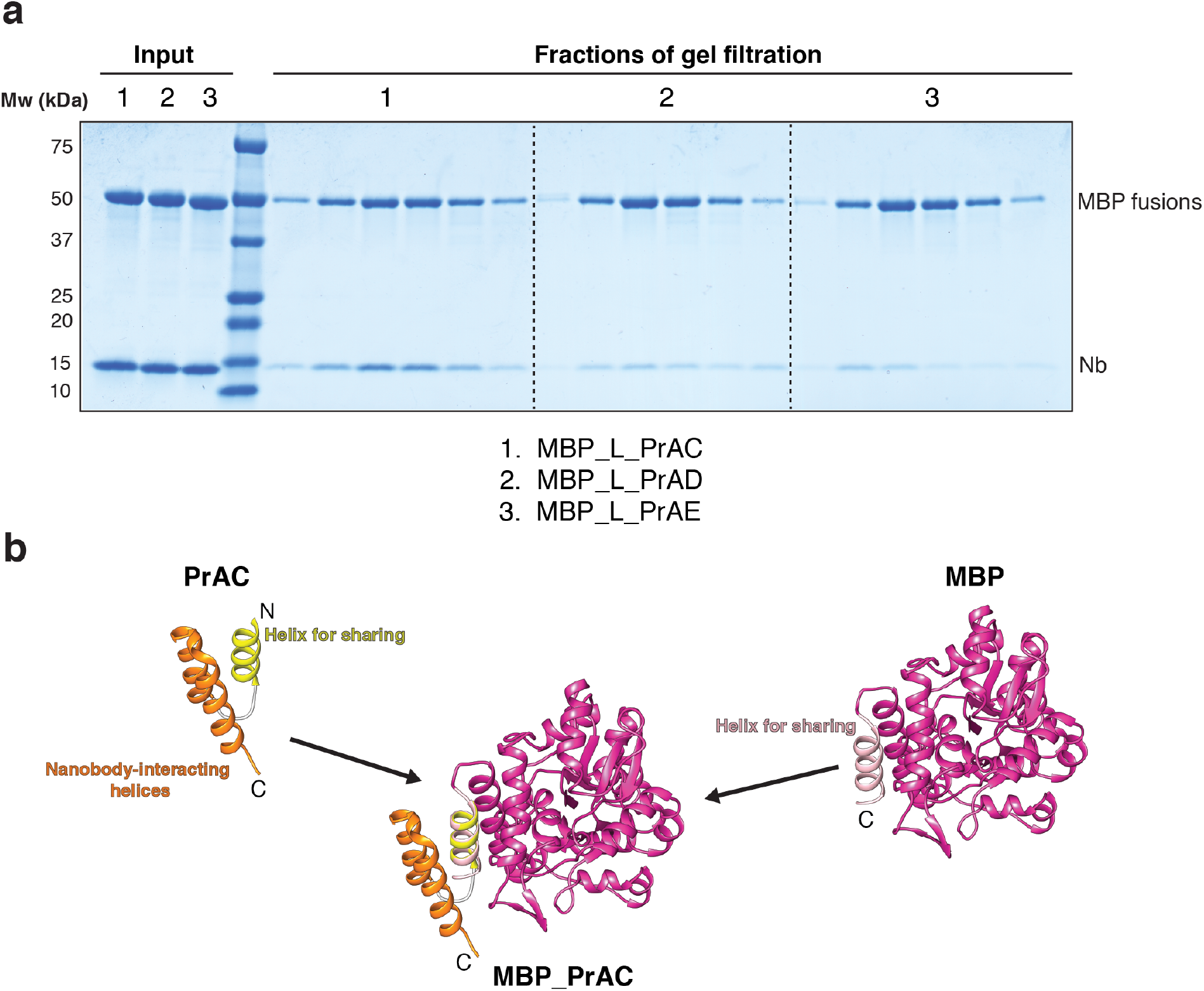
Interaction of nanobody with different PrA domains and design of a “shared helix” between PrAC and MBP. **a**, Protein A domains C, D, or E (PrAC, PrAD, PrAE) were fused to MBP via a flexible linker (L). The fusion proteins were mixed with nanobody at a 1:3 molar ratio. The samples were subjected to size-exclusion chromatography and fractions were analyzed by SDS-PAGE and Coomassie-blue staining. **b**, A “shared helix” was generated from a helix of PrAC (4NPD) that is not interacting with the nanobody and the C-terminal helix of MBP (1ANF).

**Fig. S3.**
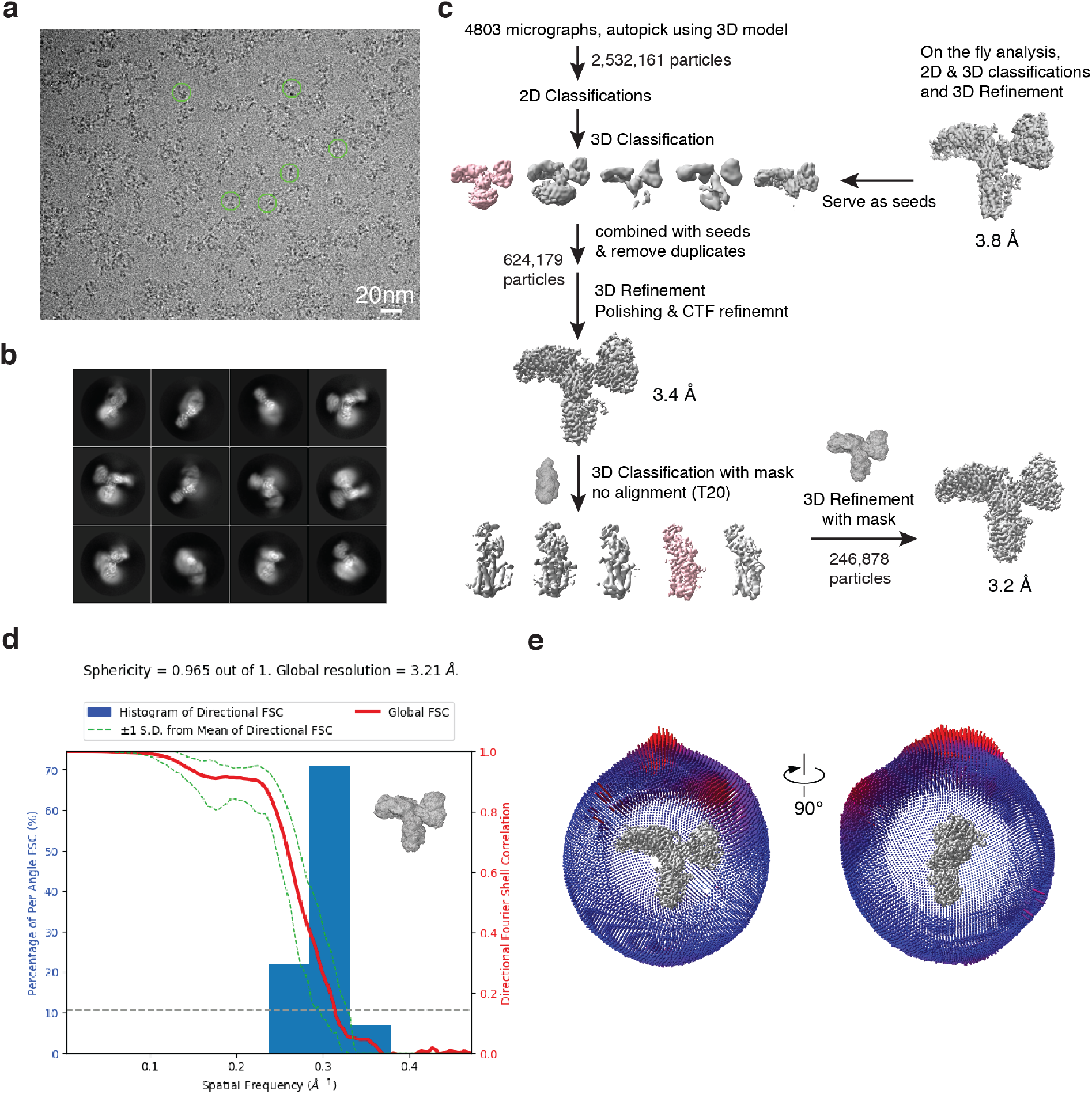
Cryo-EM analysis of the KDELR/Legobody complex. **a**, Representative cryo-EM image with selected particles marked by green circles. **b**, Representative 2D class averages of selected particles. **c**, Cryo-EM processing workflow. The classes selected for further analysis are shown in pink. Masks used for classification and refinement are indicated. **d**, 3D FSC curves and preferred orientation analysis. The red line shows the global FSC and the green dotted lines indicate the +1 and – 1 standard deviations around the Mean of Directional FSC curve. The FSC calculations used the mask shown on the side. A sphericity of 0.965 was estimated, indicating a mostly isotropic reconstruction without strong orientation bias. **e**, Euler angle distribution of refined particles is shown in two different views.

**Fig. S4.**
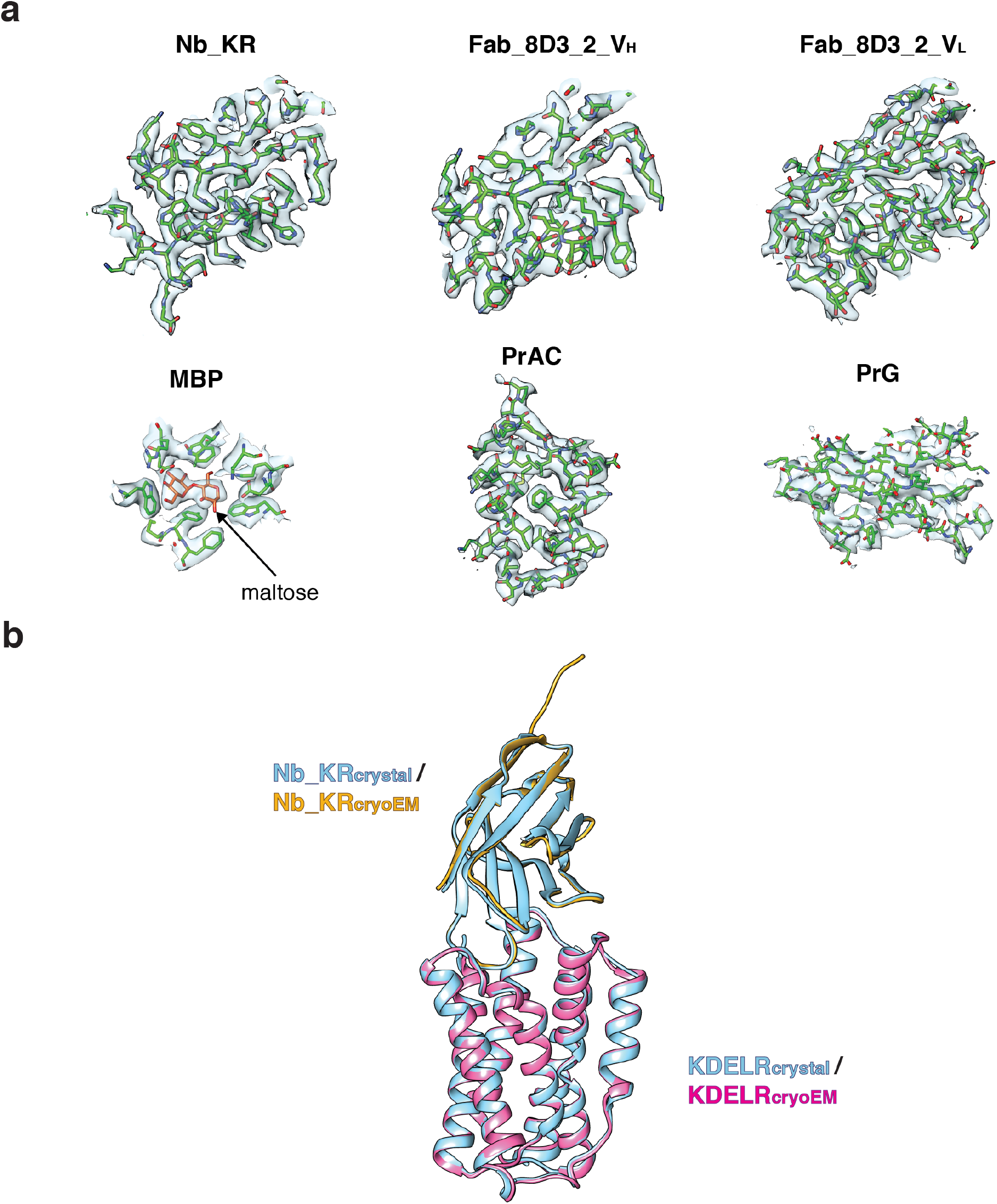
Fit of models for Legobody components and comparison of the cryo-EM and crystal structures of the KDELR/nanobody complex. **a**, Map and fitted models for different parts of the Legobody. **b**, The cryo-EM structure of the KDELR/Nb_KR complex was aligned with the structure determined by X-ray crystallography (PDB code 6I6J). The proteins are shown in cartoon representation.

**Fig. S5.**
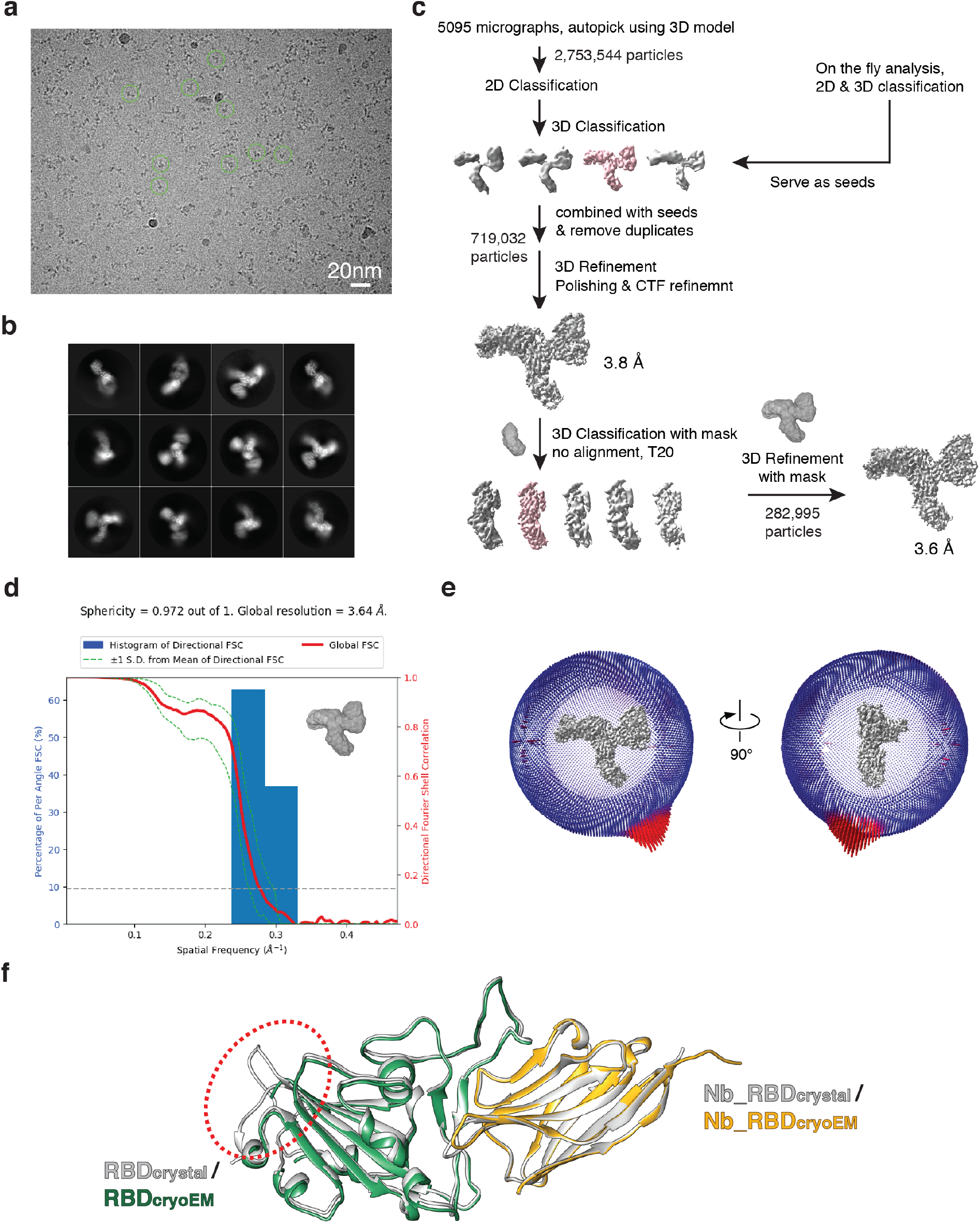
Cryo-EM analysis of the RBD/Legobody complex and comparison of the cryo-EM and crystal structures of the RBD/nanobody complex. **a**, Representative cryo-EM image with selected particles marked by green circles. **b**, Representative 2D class averages of selected particles. **c**, Cryo-EM processing workflow. The classes selected for further analysis are shown in pink. Masks used for classification and refinement are indicated. **d**, 3D FSC curves and preferred orientation analysis. The red line shows the global FSC and the green dotted lines indicate the +1 and – 1 standard deviations around the Mean of Directional FSC curve. The FSC calculations used the mask shown on the side. A sphericity of 0.972 was estimated, indicating a mostly isotropic reconstruction without strong orientation bias. **e**, Euler angle distribution of refined particles is shown in two different views. **f**, The cryo-EM structure of the RBD/Nb_RBD complex was aligned with the structure determined by X-ray crystallography (PDB code 7KGJ). The proteins are shown in cartoon representation. The red circle highlights loops that are invisible in the cryo-EM map.

**Fig. S6.**
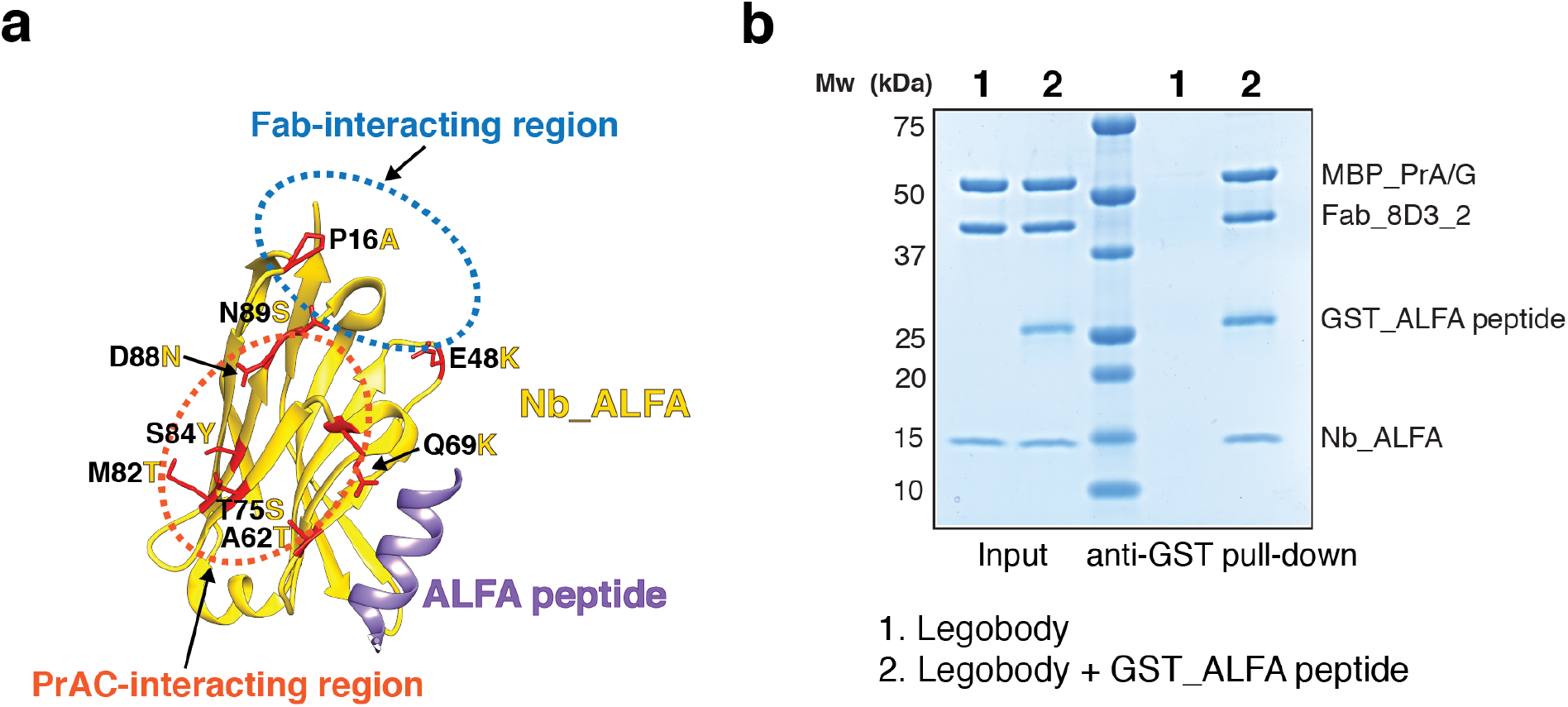
Generation of a Legobody-compatible ALFAtag nanobody. **a**, Structure of the complex of the original ALFA nanobody with the ALFA peptide. The model shows the polypeptides in cartoon representation. Residues shown as red sticks deviate from those seen in the common nanobody framework (in yellow) and would interfere with the assembly into a Legobody. They were therefore mutated to generate Nb_ALFA. **b**, To demonstrate that Nb_ALFA can be assembled into the Legobody without loss of antigen binding, purified Legobody containing Nb_ALFA was incubated with or without a GST fusion of ALFA peptide (GST_ALFA peptide). The samples were incubated with glutathione beads and the bound material analyzed by SDS-PAGE and Coomassie-blue staining. The input corresponds to 50% of the material used for the pull-down.

**Table S1.** Sequences of all Proteins. Please contact authors directly.

**Table S2.** X-Ray data collection, and refinement statistics.

|  | Nb_0/Fab_8D3 |
| --- | --- |
| **Data Accession** |  |
| PDB | xxxx |
| **Data collection** |  |
| Space group | P 1 21 1 |
| Cell dimensions |  |
| *a*, *b*, *c* (Å) | 64.51, 60.287, 77.244 |
| $\alpha$, $\beta$, $\gamma$ (°) | 90.00, 109.341, 90.00 |
| Resolution (Å) | 1.83 (1.895-1.83) * |
| *R*<sub>merge</sub> | 0.068 (0.55) |
| *I* / σ*I* | 16.46 (2.95) |
| Completeness (%) | 99.35 (99.15) |
| Redundancy | 7.6 (7.1) |
| **Model Refinement** |  |
| Software | Phenix |
| Resolution (Å) | 1.83 |
| No. Reflections | 49251 |
| *R*<sub>work</sub> / *R*<sub>free</sub> | 0.18/0.21 |
| Number of non-hydrogen atoms | 4705 |
| Protein | 4211 |
| Water | 494 |
| Protein Residues | 554 |
| Average B factors (Å<sup>2</sup>) |  |
| Protein | 36.13 |
| R.m.s. deviations |  |
| Bond lengths (Å) | 0.01 |
| Bond angles (°) | 0.91 |
| Ramachandran statistics (%) |  |
| Outliers | 0 |
| Allowed | 2.38 |
| Favored | 97.62 |
| MolProbity score | 1.37 |
| All-atom clashscore | 5.43 |
| Poor rotamers (%) | 0.64 |

\*Values in parentheses are for highest-resolution shell.

**Table S3.** Cryo-EM data collection, refinement, and validation statistics.

| Structure | KDEL<sup>R</sup>/<br>Legobody | RBD/<br>Legobody |
| --- | --- | --- |
| **Data Accession** |  |  |
| PDB | xxxx | xxxx |
| EMDB | EMD-xxxxx | EMD-xxxxx |
| **Data Collection** |  |  |
| Microscope | Titan Krios | Titan Krios |
| Voltage (kV) | 300 | 300 |
| Exposure navigation | Stage Movement<br>&beamshift | Stage Movement<br>&beamshift |
| Automation software | SerialEM | SerialEM |
| Detector | Gatan K3 (Counting) | Gatan K3 (Counting) |
| Energy filter | 25 eV | 25 eV |
| Nominal magnification | 81k | 81k |
| Pixel Size (Å/pixel) | 1.06 | 1.06 |
| Exposure time (s), frames | 2.2, 50 | 2.16, 50 |
| Exposure rate (e<sup>-</sup> pixel<sup>-1</sup> s<sup>-1</sup>) | 26.31 | 26.18 |
| Electron exposure (e<sup>-</sup>/Å<sup>2</sup>) | 51.44 | 50.42 |
| Defocus range (µm) | -1.0 to -2.6 | -1.0 to -2.6 |
| Micrographs collected | 4,863 | 5795 |
| **Reconstruction** |  |  |
| Software | Relion 3.1 | Relion 3.1 |
| Micrographs used | 4,803 | 5095 |
| Particles used in refinement | 246,878 | 282,995 |
| Symmetry | C1 | C1 |
| Overall resolution (Å)<br>FSC=0.143 (masked) | 3.2 | 3.6 |
| Map sharpening B-factor (Å<sup>2</sup>) | -100 | -130 |
| Local resolution range (Å) | 3.0 ~ 5.0 | 3.4 ~ 5.4 |
| **Model Refinement** |  |  |
| Software | Phenix | Phenix |
| Non-hydrogen atoms | 9361 | 9008 |
| Protein residues | 1178 | 1148 |
| Ligands | 5 | 1 |
| Average B factors (Å<sup>2</sup>) |  |  |
| Protein | 42.4 | 34.64 |
| Ligands | 34.2 | 34.45 |
| R.M.S. deviations |  |  |
| Bond length (Å) | 0.003 | 0.003 |
| Bond angle (°) | 0.578 | 0.615 |
| Ramachandran statistics (%) |  |  |
| Outliers | 0 | 0 |
| Allowed | 1.91 | 2.59 |
| Favored | 98.09 | 97.41 |
| MolProbity score | 1.45 | 1.64 |
| Clashscore | 8.22 | 10.03 |
| Poor rotamers (%) | 0 | 0 |
| Model vs. Map FSC |  |  |
| FSC=0.5 (masked, Å) | 3.3 | 3.8 |

**Table S4.** Analysis of clashes between Legobody domains and targets. Crystal structures of target/nanobody complexes were aligned with the structure of the Legobody on the basis of the nanobody. Steric clashes with the different Legobody domains are listed. The Table shows only one entry for a target protein if several similar structures have been reported. In some cases, the existences of clashes are uncertain (labeled as “possible”).

**Small membrane proteins (<100 kDa)**
| Target Name | PDB code | No Clash | Clash with PrAD | Clash with MBP_PrAC | Clash with Fab_8D3 |
| --- | --- | --- | --- | --- | --- |
| beta2 adrenoceptor | 3P0G | X |  |  |  |
| M2 muscarinic acetylcholine receptor | 4MQS | X |  |  |  |
| succinate receptor | 6IBB |  | Possible |  |  |
| Viral GPCR US28 | 5WB1 | X |  |  |  |
| opioid receptor | 5C1M |  |  | X |  |
| KDEL receptor | 6I6J | X |  |  |  |
| lactose permease | 5GXB | X |  |  |  |
| L-amino acid transporter | 6F2G |  | X |  |  |
| ADP/ATP Carrier | 6GCI | X |  |  |  |
| dipeptide and tripeptide permease | 6GS4 | X |  |  |  |
| glycine transporter | 6ZBV | X |  |  |  |
| divalent metal ion transporter | 5M94 |  |  | X |  |
| MraY translocase | 6OYH | X |  |  |  |
| insertase BamA | 6QGX | X |  |  |  |

**Small soluble Proteins (<100 kDa)**
| **Target Name** | **PDB code** | **No Clash** | **Clash with PrAD** | **Clash with MBP_PrAC** | **Clash with Fab_8D3** |
| --- | --- | --- | --- | --- | --- |
| RBD of SARS-CoV-2 | 7KGJ | X |  |  |  |
| ALFA | 6I2G | X |  |  |  |
| GFP | 3K1K | X |  |  |  |
| BC2 peptide tag | 5IVN | X |  |  |  |
| D3 domain of MTIP | 4GFT | X |  |  |  |
| Complement C3 | 6XZU | X |  |  |  |
| Vsig4 | 5IML | X |  |  |  |
| PorM | 6EY0 | X |  |  |  |
| Lysozyme | 3EBA | X |  |  |  |
| Nucleoporin-98 | 7NOW |  |  | Possible |  |
| Gelsolin | 6H1F | X |  |  |  |
| CD38 | 5F1O | X |  |  |  |
| fimbrial adhesin AC | 4W6X | X |  |  |  |
| Surface protein P12p | 7KJI | X |  |  |  |
| MAGEB1 | 6R7T | X |  |  |  |
| Capsid protein of norovirus | 5O02 | X |  |  |  |
| Nucleoporin NIC96 | 6X07 |  |  | X |  |
| Complement C1q | 6Z6V | X |  |  |  |
| PD-L1 | 5JDS | X |  |  |  |
| CTLA-4 | 6RQM | X |  |  |  |
| Nup120 | 6X06 | X |  |  |  |
| DHFR | 4EIZ | X |  |  |  |
| GAK kinase | 4C59 | X |  |  |  |
| Synaptojanin1 | 7A17 | X |  |  |  |
| Human prion protein | 4N9O | X |  |  |  |
| CD9 | 6Z1V | X |  |  |  |
| Nup54 | 5C2U | X |  |  |  |
| Urokinase-type plasminogen activator | 5LHP | X |  |  |  |
| EDS1 | 6I8H | X |  |  |  |
| Octarellin V.1 | 5BOP | X |  |  |  |
| Capsid protein p24 | 5O2U | X |  |  |  |
| Ricin | 4LGP | X |  |  |  |
| Cyclin-G-associated kinase | 5Y7Z | X |  |  |  |
| CD1d | 6V7Y | X |  |  |  |
| Toxin HigB-2 | 5JA8 | X |  |  |  |
| RhoB | 6SGE | X |  |  |  |
| human serum albumin | 5VNW |  |  |  | X |

